# Phosphoproteomic analysis reveals the diversity of signaling behind ErbB inhibitor-induced phenotypes

**DOI:** 10.1101/2024.03.12.584564

**Authors:** Katri Vaparanta, Anne Jokilammi, Johannes Merilahti, Johanna Örling, Noora Virtanen, Cecilia Sahlgren, Klaus Elenius, Ilkka Paatero

## Abstract

The impact of kinase inhibitors on the phosphoproteome has been rarely investigated at a whole organism level. Here we performed a phosphoproteomic analysis in embryonic zebrafish to identify the signaling pathways perturbed by ErbB receptor tyrosine kinase inhibitors at the organism level. The phosphorylation of proteins associated with the PI3K/Akt, p38 MAPK, Notch, Hippo/Yap and β-catenin signaling pathways were differentially regulated by the ErbB inhibitors. Gene set enrichment analyses indicated differential neurological and myocardial phenotypes of different ErbB inhibitors. To assess the neurological and myocardial effects, motility and ventricle growth assays were performed on zebrafish embryos treated with the ErbB and downstream signaling pathway inhibitors. The treatment with the inhibitors targeting the PI3K/Akt, p38 MAPK, and Notch signaling pathways along with the ErbB inhibitors AG1478 and Lapatinib perturbed the overall movement and ventricle wall growth of zebrafish embryos. Taken together, these results indicate that inhibitors with the same primary targets can affect different signaling pathways while eliciting similar physiological phenotypes.

## Introduction

Several molecularly targeted therapies have been developed to treat cancer and many of these agents target cellular signaling systems and pathways (Min & Lee, 2022). One of the first signaling systems to be targeted was ErbB/Her receptor tyrosine kinases (Min & Lee, 2022). Several antibodies and small molecules have been developed to target ErbB signaling with differential effects in clinical trials (Roskoski, 2019), indicating differences in the mechanism of action and/or toxicity profiles.

Interestingly, compounds with similar primary targets *in vitro* may have quite different effects *in vivo*. This indicates that *in vivo* analysis of the effects of compounds on cell signaling could be used to elucidate the organismal level changes in their actions. This is, however, difficult to achieve by using mammalian *in vivo* models or human samples, as already the size and complexity hinder effective whole-organism analyses of signaling effects. In recent years, zebrafish embryos have been utilized increasingly as an *in vivo* model to analyze the *in vivo* effects of compounds (MacRae & Peterson, 2015). The small size and large number of zebrafish embryos enable analysis of changes of signaling at the whole organism level for example using phosphoprotemic analysis (Lemeer et al, 2008; Kwon et al, 2016) without the need for prior selection of tissues or cell types of interest. Mass-spectrometric analysis of phosphoproteome has been, however, underutilized in studies of the effects of chemical compounds on zebrafish development and signaling.

Phenotypic analyses of zebrafish embryos exposed to ErbB inhibitors lapatinib, gefitinib, or AG1478 have shown differential phenotypes in cardiac biology and in the overall movement of the zebrafish embryos (Paatero et al, 2019; Vaparanta et al, 2023). Both the ventricle wall growth and the overall movement of the zebrafish embryos have been most strongly affected by AG1478 treatment and least with gefitinib treatment (Paatero et al, 2019; Vaparanta et al, 2023). This prompted us to analyze the signaling differences of these inhibitors in-depth using analysis of whole organism phosphoproteome using protein mass spectrometry.

## Results

To explore the global phosphoproteomic changes induced by the ErbB kinase inhibitors known to elicit differential phenotypic effects, we exposed developing zebrafish embryos to lapatinib, gefitinib, AG1478 or DMSO (Fig. 1A). The phosphopeptides from the embryos were enriched and analyzed with protein mass spectrometry using data-independent acquisition with MS/MS (Gerritsen & White, 2021; Higgins et al, 2023). In total 23,141 unique phosphopeptides were detected (Supplemental Table 1) and the quantities of 2033 phosphopeptides were significantly altered between treatments (Supplemental Table 2). Differential expression analysis on the phosphorylated peptides in each condition was conducted and hierarchical clustering was utilized to identify differentially regulated clusters of phosphopeptides. Even though the inhibitors are structurally related (Fig. 1B), clear differences in the size of the phosphopeptide clusters were observed (Fig. 1C). An unexpectedly large cluster of phosphopeptides that were increased in lapatinib-treated embryos but reduced in AG1478 and gefitinib treated embryos was detected (Cluster 1 in Fig. 1C). Overrepresentation analysis indicated that phosphopeptides in this cluster were associated with the p38 MAPK, ERK and mTORC1 pathways (Fig. 1C). The clusters of phosphopeptides that were increased only in either AG1478 (Cluster 3 in Fig. 1C) or gefitinib (Cluster 6 in Fig. 1C) treated embryos but reduced in embryos treated with the other inhibitors were in turn significantly smaller.

**Figure 1.**
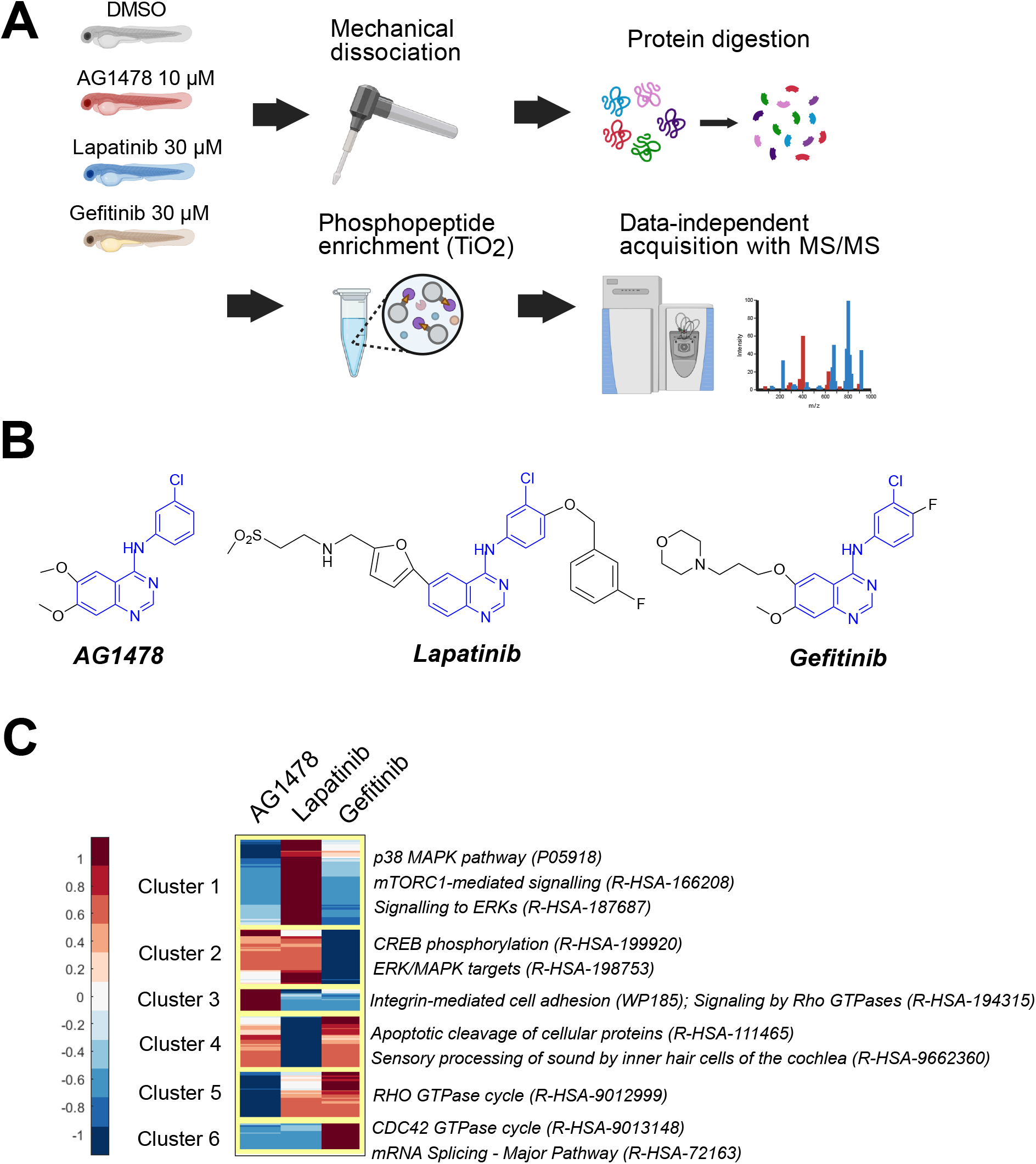
Whole organism phosphoproteome of ErbB inhibitor treated zebrafish embryos. A: The workflow for the global phosphoproteome acquirement from ErbB inhibitor-treated zebrafish embryos. B: The chemical structure of the ErbB inhibitors. The shared core structure is indicated in blue. C: Heatmap of the differently regulated phosphopeptides. The median fold change against the control DMSO treatment of four replicate experiments is visualized on a standardized scale. The clusters were identified with hierarchical clustering. Overrepresentation testing was used to associate pathways for each cluster.

The differentially regulated individual phosphorylated peptides in each condition were visualized in scatter plots (Fig. 2A-C). In AG1478 and gefitinib-treated embryos, the downregulation of the phosphorylation of the activating residues of ERK MAPKs (Mapk1/3) and p38 MAPKs (Mapk11/12/13/14) was observed along with the phosphorylation of S6 kinases (Rps6ks) and other substrates (Akt1s1) downstream of the Akt/mTOR pathway. In the lapatinib-treated embryos instead, the phosphorylation of both p38 MAPKs and S6 kinases was upregulated. In the lapatinib-treated embryos, the phosphorylation of proteins involved in the Hippo/YAP pathway (Lats1), transcriptional repressors of Notch signaling (Ncor1/2, Snw1), and transcriptional activators of β-catenin (Bcl9/Bcl9l) were downregulated. In gefitinib-treated embryos, however, the phosphorylation of the transcriptional repressors of Notch signaling (Hdac1, Spen, Ncor1) was upregulated. In AG1478-treated embryos, in turn, the phosphorylation of the transcriptional activators of β-catenin (Bcl9/Bcl9l) was significantly increased. Gene set enrichment analyses (GSEAs) (Subramanian et al, 2005) were performed to identify the dysregulated signaling pathways from the whole dataset. The results from the GSEAs confirmed the results from significantly differentially regulated phosphorylated residues (Fig. 2D-F, supplemental table 3). The phosphorylation events associated with the p38 MAPK and PI3K/Akt/mTOR signaling pathways were discovered to be upregulated by lapatinib and downregulated by AG1478 and gefitinib treatment (Fig. 2D-F). The phosphorylation of proteins involved in ERK MAPK signaling was downregulated by gefitinib and AG1478 treatment (Fig. 2D,F) while the phosphorylation of proteins involved in Hippo/YAP, Notch, and transcriptional β-catenin signaling was downregulated by lapatinib treatment only (Fig. 2E). Taken together, these results suggest that multiple signaling pathways are differently regulated by the gefitinib, lapatinib and AG1478 treatment.

**Figure 2.**
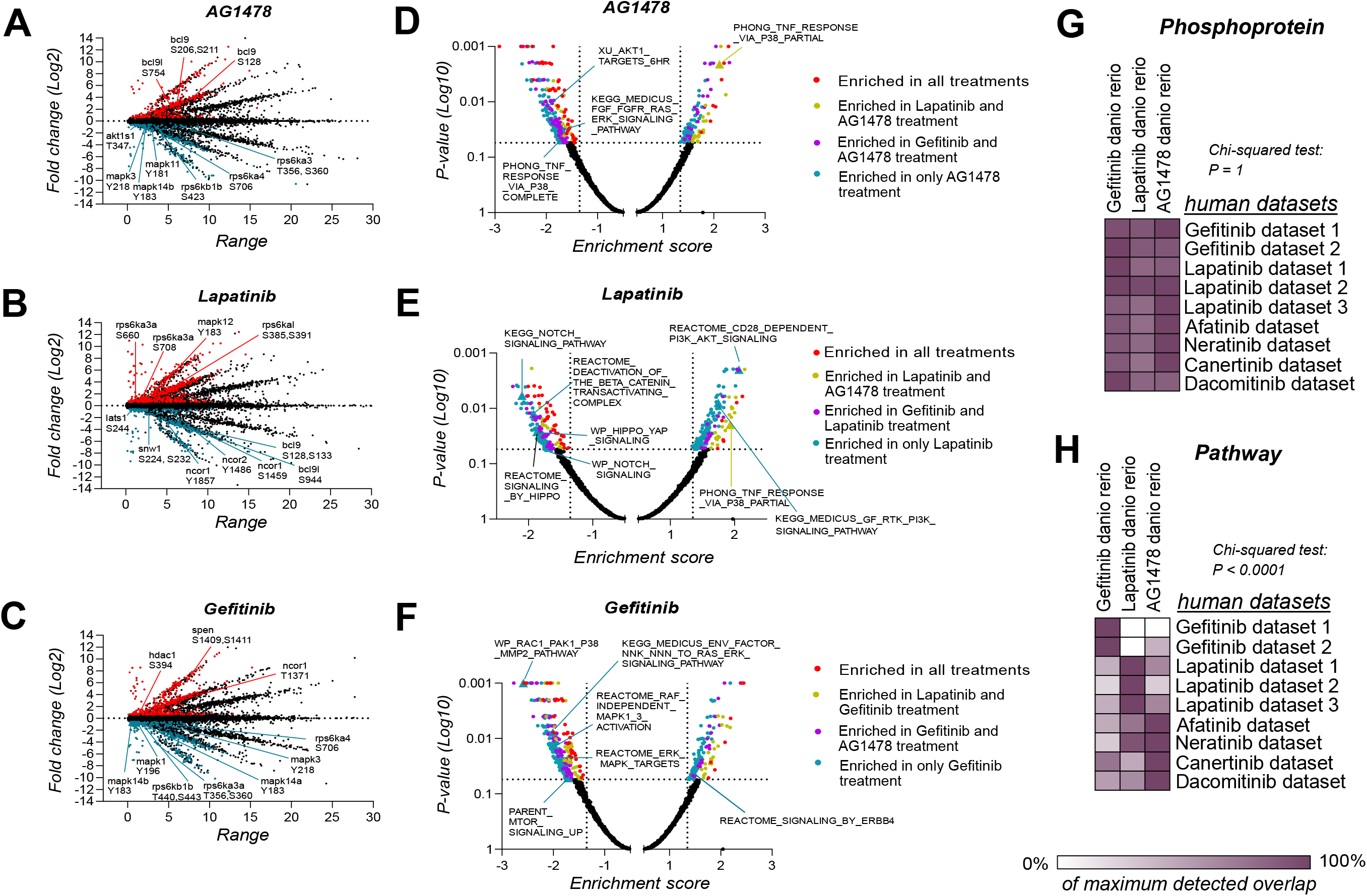
Phosphorylation of proteins involved in the Notch, Erk MAPK, PI3K/Akt/mTOR, p38 MAPK and β-catenin signaling pathways is differently regulated by the ErbB inhibitors. **A-C:** Scatter plots of the identified phosphopeptides from the MS/MS analysis in each condition. The significantly downregulated (blue) and upregulated (red) phosphopeptides are indicated. Median fold change against the DMSO control treatment and the value range of four replicate experiments in log2 scale are visualized in the y- and x-axis, respectively. **D-F**: Volcano plots of the results of the gene set enrichment analysis performed on the global phosphoproteomics data. **G**: Overlap of the differentially phosphorylated proteins between the MS/MS analysis results and previously acquired human datasets of the ErbB inhibitor-regulated phosphoproteome. **H**: Overlap of the overrepresented pathways of the differentially phosphorylated proteins between the MS/MS analysis results and previously acquired human datasets of ErbB inhibitor-regulated phosphoproteome.

The overlap of the results of the phosphoproteomic analysis in zebrafish with published datasets from ErbB inhibitor treated human samples was explored both at the phosphoprotein and signaling pathway level. Analysis on the phosphoprotein level indicated only limited specificity of overlap between different ErbB inhibitors (Fig. 2G). Pronounced differences in the overlap of signaling pathways, however, were discovered between ErbB inhibitor treatments (Fig. 2H). Significantly more overlap between the signaling pathways of zebrafish samples treated with gefitinib with the human samples treated with gefitinib than with the human samples treated with lapatinib or with the pan-ErbB inhibitors (afatinib, neratinib, canertinib, dacomitinib), was observed. Similarly, the signaling pathways perturbed in zebrafish embryos by lapatinib treatment or AG1478 treatment overlapped most with the signaling pathways perturbed in the human samples treated with lapatinib or pan-ErbB inhibitors, respectively. This indicates that the signaling differences observed in the human samples treated with these inhibitors are recapitulated in the zebrafish context on the signaling pathway level.

To validate the findings of the phosphoproteomic analysis, the phosphorylation of the two activating residues of p38 MAPKs was measured from gefitinib, lapatinib, AG1478 and control-treated zebrafish embryos with Western analyses. As expected from the results of the phosphoproteomic analysis, the overall (Fig. 3A) or simultaneous double (Fig. 3B) phosphorylation of the activating residues of p38 MAPKs was increased by lapatinib treatment and decreased by gefitinib treatment compared to other treatments (Fig. 3A-B). To examine the correlation of previously (Vaparanta et al, 2023) and newly executed Western analyses with the results of the phosphoproteomic analysis, correlation analyses were performed. The intensity values of phosphosite-specific antibodies in Western analyses had a significant positive correlation with the intensity values of the phosphopeptides identified in the phosphoproteomic analysis and the enrichment scores of signaling pathways from the GSEAs that were performed on the phosphoproteomics data. This suggests that the phosphoproteomic analysis reliably measured signaling activities in zebrafish embryos.

**Figure 3.**
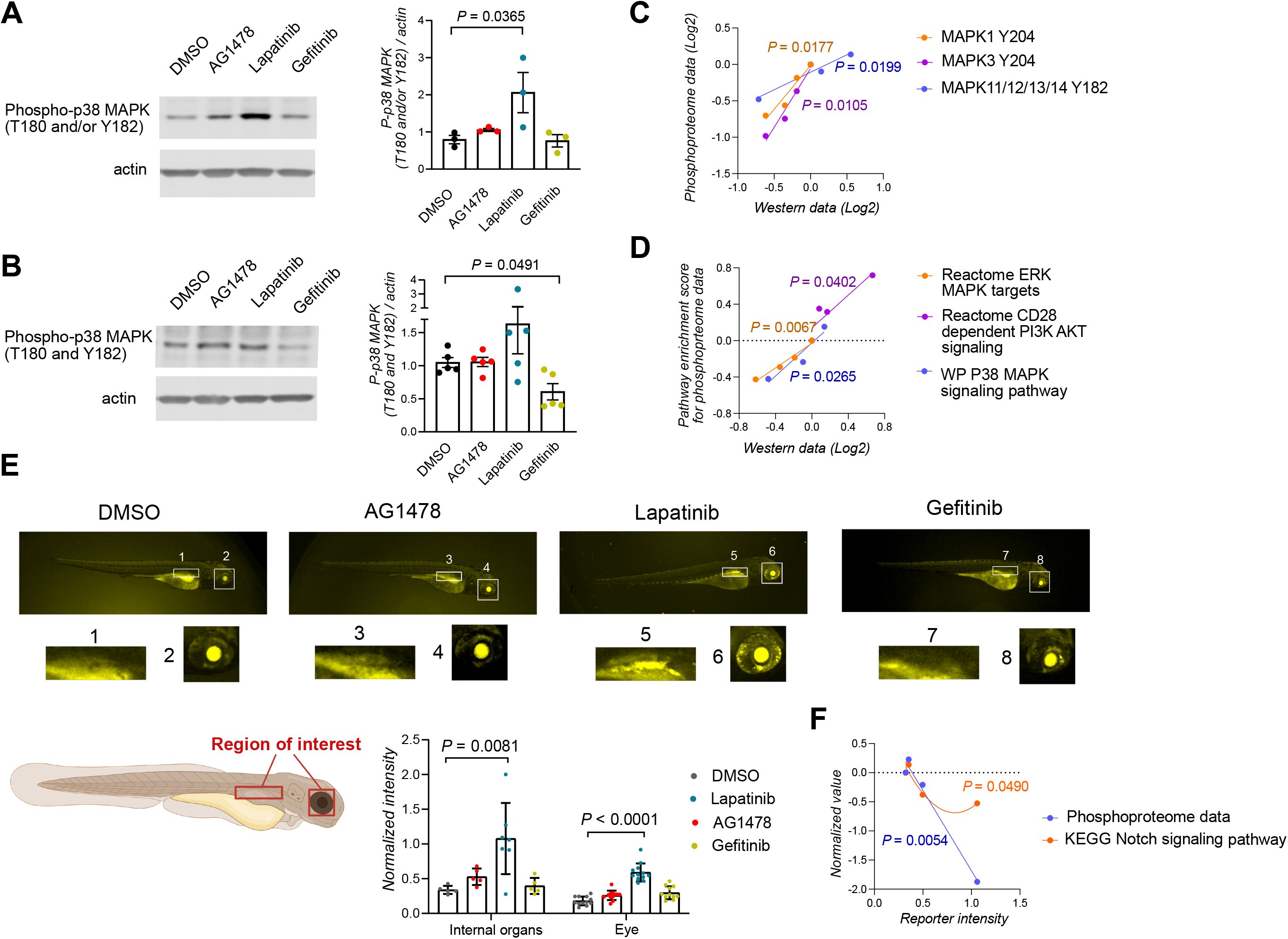
Validation of the ErbB inhibitor-induced signaling changes identified through the whole organism phosphoproteome analysis. **A-B:** Representative blots and densitometric quantification of the western analyses on the activation of p38 MAPK kinases in ErbB inhibitor-treated zebrafish embryos at 3dpf. Mean ±SD. One dot represents one replicate experiment. One-way ANOVA (A) and Welch’s ANOVA (B) were used for statistical testing. Dunnett’s multicomparison test was utilized for ANOVA post hoc analyses. **C-D:** Correlation of previously acquired (Vaparanta et al, 2023) and new Western analyses with the global phosphoproteome data at the phosphosite level (C) and with the GSEA enrichment scores of the pathways enriched in the phosphoproteome data (D). The mean LFQ-intensity values of the phosphopeptides containing the phosphorylated residue and densitometric measurements of the western analyses were normalized against the DMSO control and transformed to the Log2 scale. One-tailed Pearson correlation analysis was utilized. One dot represents the value of one treatment condition (DMSO, AG1478, Lapatinib, Gefitinib) in the Western and MS/MS or gene set enrichment analyses. **E:** Fluorescence microscope images and intensity measurements of the tp1:Venus-pest Notch reporter line (Ninov et al, 2012) zebrafish embryos treated with the indicated ErbB inhibitors or DMSO at 3 dpf. The regions of interests have been outlined in the schematic. Kruskal Wallis and Dunn’s multicomparison test was utilized for statistical analysis. One dot corresponds to one zebrafish embryo. Mean ±SD. **F:** Correlation and nonlinear relationship analysis of the mean LFQ intensity of the phosphopeptides of proteins involved in the Notch signaling pathway or the GSEA enrichment scores and the Notch reporter fluorescence intensity values. One-tailed Pearson correlation and least squares fitting were used for the analyses. The statistical significance of the R^2^ for the quadratic fit was calculated with an F-test. One dot represents the value of one treatment condition (DMSO, AG1478, Lapatinib, Gefitinib) in the Notch reporter and MS/MS or gene set enrichment analyses.

For additional validation, the effect of ErbB inhibitor treatment on the transcriptional activity of Notch was measured by utilizing a transgenic Notch activity reporter zebrafish line (tp1:VENUS-PEST) (Ninov et al, 2012). The signal from the reporter has been detected in several tissues in the zebrafish embryos such as the eye (MacDonald et al, 2015) and several internal organs such as the pancreas at 2-4 dpf (Ghaye et al, 2015) and in the liver at 4 dpf (So et al, 2018). The fluorescence signal from the Notch reporter was observed to be more intense in the eye and internal organs in the lapatinib-treated embryos compared to gefitinib, AG1478, or control-treated embryos (Fig. 3E). This suggests that the reduced phosphorylation of the transcriptional repressors of Notch signaling by lapatinib treatment increased the transcriptional activity of Notch in the embryos. The relationship of the observed fluorescent signal with the intensity values of the phosphopeptides of transcriptional Notch repressors identified in the phosphoproteomic analysis and the enrichment score of signaling pathways from the GSEAs that were performed on the phosphoproteomics data was examined. A negative correlation and a nonlinear relationship were discovered between the Notch reporter intensity values and the phosphopeptide intensities and the enrichment score of the KEGG Notch signaling pathway, respectively (Fig. 3F). This further suggests that the discoveries from the phosphoproteomic analysis are reproducible by other methodologies.

A GSEA of the ontologies of the human phenotype ontology database (Köhler et al, 2021) was performed on the phosphoproteomic data (Fig. 4A). Differences in the phenotypes associated with the phosphorylation changes induced by the ErbB inhibitors were observed. As also previously reported (Paatero et al, 2019; Vaparanta et al, 2023), myocardial and neurological phenotypes were only associated with AG1478 and lapatinib treatment. Since the phosphoproteomic changes induced by the treatment of these inhibitors were markedly different, we wanted to explore if blocking the pathways differentially regulated by AG1478 and lapatinib would result in similar neurological and myocardial phenotypes in embryonic zebrafish to AG1478 or lapatinib treatment. To analyze the myocardial phenotype, zebrafish embryos were treated for 2 days with the ErbB inhibitors and inhibitors targeting the PI3K/Akt (LY294002), p38 MAPK (SB38053 and BIRB796), Notch (DAPT), and Hippo/YAP (VT103) pathways. The ventricle wall area was measured with in vivo imaging of zebrafish embryo hearts at 4dpf (Fig. 4B). A distinct difference in the ventricle wall growth was detected by the treatment of inhibitors targeting the PI3K/Akt, p38 MAPK, and Notch signaling along with AG1478 and lapatinib treatment (Fig. 4C). To analyze the differential effects on neurological phenotypes suggested by the GSEA (Fig. 4A), we exposed zebrafish embryos to the inhibitors and carried out motility assays (Fig. 4D). The inhibitors targeting PI3K/Akt, p38 MAPK, and Notch signaling affected the motility of the zebrafish embryos along with AG1478 and lapatinib (Fig. 4E). This along with the results of the phosphoproteomic analysis suggests that these differentially regulated pathways are involved in the phenotypic changes induced by AG1478 and lapatinib treatment.

**Figure 4.**
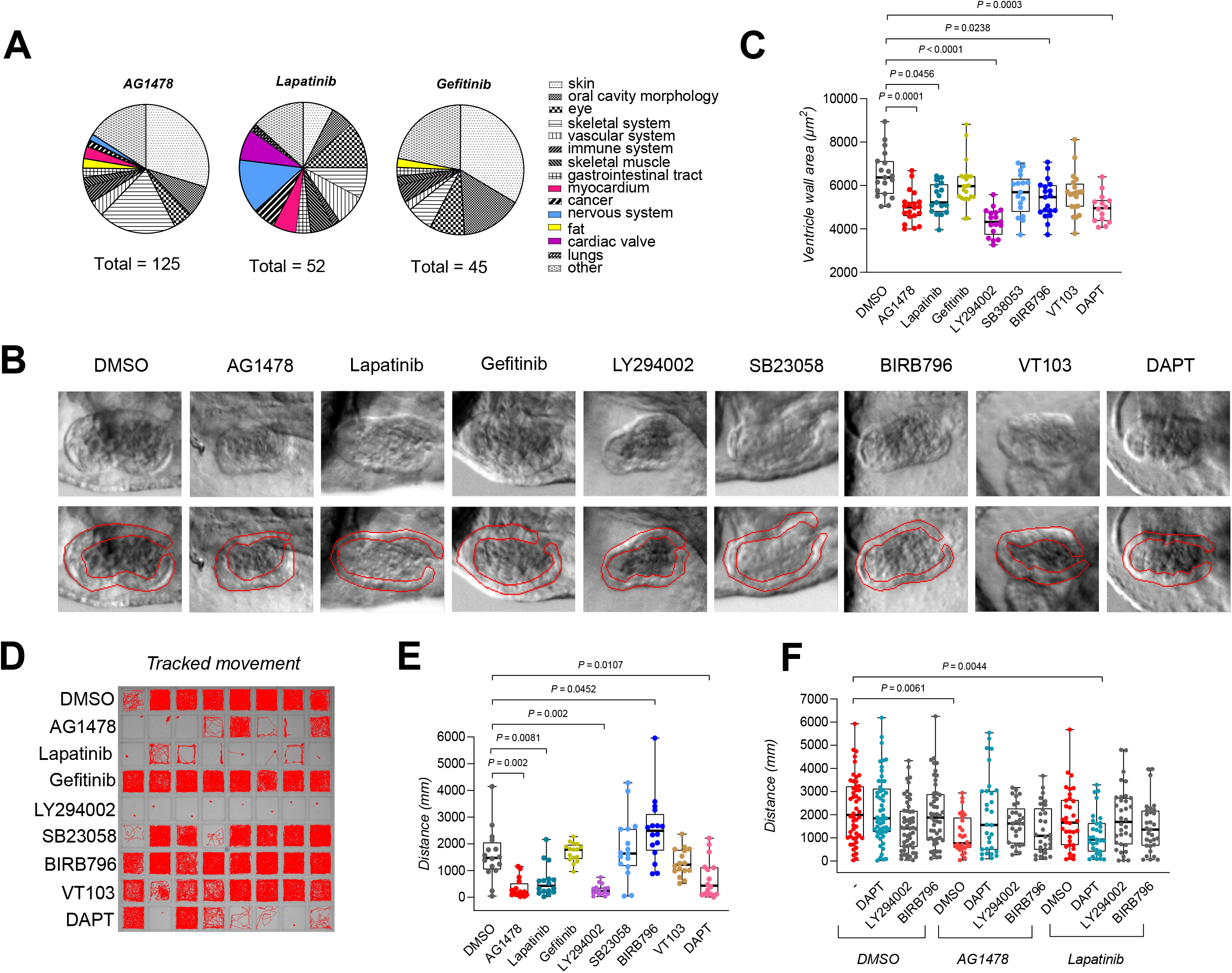
Phenotypic changes induced by inhibitors targeting the ErbB receptors and the signaling pathways differentially perturbed by the ErbB inhibitors. A: Enriched phenotypes of ErbB inhibitor-treated zebrafish embryos according to the GSEA on the global phosphoproteomics data. Only negatively enriched phenotypes were included and categorized into visualized subcategories. B-C: Phase-contrast images and quantification of the ventricle wall thickness of control or inhibitor-treated zebrafish embryos at 4dpf. Inhibitors targeting the ErbB receptors (AG1478 (10μM), lapatinib (30μM), gefitinib (30μM), the PI3K/Akt pathway (LY294002, 30μM), the p38 MAPK pathway (SB23058, 30μM, BIRB796, 30μM), the Hippo/YAP pathway (VT103, 30μM) and the Notch pathway (DAPT, 50μM) were examined. Kruskal-Wallis ANOVA and Dunnett’s multicomparison test were used for statistical testing. One dot corresponds to one zebrafish embryo. In the boxplots, the line denotes the median, the box the interquartile range, and whiskers the whole range of values. D-E: Motility of control or inhibitor-treated zebrafish embryos at 4dpf. Inhibitors targeting the ErbB receptors (AG1478 (10μM), lapatinib (30μM), gefitinib (30μM)), the PI3K/Akt pathway (LY294002, 30μM), the p38 MAPK pathway (SB23058, 30μM, BIRB796, 30μM), the Hippo/YAP pathway (VT103, 30μM) and the Notch pathway (DAPT, 50μM) were analyzed. The tracked path (D) and the quantifications of the traveled distance are visualized (E). Kruskal-Wallis ANOVA and Dunnett’s multicomparison test were used for statistical testing. One dot corresponds to one zebrafish embryo. In the boxplots, the line denotes the median, the box the interquartile range and whiskers the whole range of values. F: Motility of co-treated zebrafish embryos at 4dpf. Embryos treated with the inhibitors targeting the PI3K/Akt (LY294002), the p38 MAPK (BIRB796), Notch (DAPT) and ErbB (AG1478, lapatinib) pathways were examined alone or in combination in low doses. The quantifications of the traveled distance are visualized. Kruskal-Wallis ANOVA and Dunn’s multicomparison test were used for statistical testing. One dot corresponds to one zebrafish embryo. In the boxplots, the line denotes the median, the box the interquartile range, and whiskers the whole range of values.

To further explore this hypothesis, the effect of the co-treatment of AG1478 or lapatinib with inhibitors targeting either the PI3K/Akt (LY294002), p38 MAPK (BIRB796), or Notch (DAPT) signaling pathway was analyzed. We hypothesized that the involvement of different pathways would affect sensitivity for different inhibitors. This hypothesis was tested by using low concentrations of inhibitors alone or as co-treatments. The co-treatment of lapatinib with DAPT reduced the motility of the embryos beyond that of single-agent treatment, whereas this effect was not seen in co-treatment with AG1478 and DAPT (Fig. 4F). Taken together, these results suggest that AG1478 and lapatinib may induce similar phenotypes by partially perturbing different signaling pathways in zebrafish embryos.

## Discussion

We carried out in-depth phosphoproteomic analysis of three structurally related ErbB inhibitors (AG1478, lapatinib, gefitinib) at the whole-organism level on zebrafish embryos. The effects of small molecule inhibitors on phosphoproteome have been rarely observed at a whole organism level although these compounds are known to affect the organism globally. More targeted phosphoproteomic analyses may be unable to detect all perturbed pathways since the perturbation might be limited to unprobed organs or arise from the intercommunication between tissues.

The ErbB inhibitors we chose to investigate all share the same core structure, which targets the ATP binding pocket of the kinase domain of ErbB receptors (Roskoski, 2019). Despite these similarities, their effects on the global phosphoproteome were significantly different. These differences in effects on downstream signaling may explain also the observed differential biological effects (Paatero et al, 2019; Vaparanta et al, 2023). ErbB inhibitors differ in their efficacy and toxicity also in clinical setting (Dai et al, 2021), and one underlying cause could be differential effects on cell signaling. The global phosphoproteomic approach we employed here could be more widely utilized to analyze the differential signaling effects of closely related compounds. Especially the pathways for which no specific tools exist could be probed in this manner. This interpretation is supported by our experiments where we observed several differentially affected pathways that do not have these specific tools necessary available. With our approach, we were able to link the p38 MAPK, Notch, and PI3K/Akt-signaling pathways into ErbB-inhibitor-induced neurological and myocardial phenotypes.

We utilized the results of the global phosphoproteomic analysis further by carrying out extensive analyses to identify altered pathways. Indeed, we were able to confirm several identified pathways using phospho-specific antibodies. These antibodies exist only for a limited number of potential targets, and antibodies suited for work with zebrafish material are even more scarce (Paardekooper Overman & Den Hertog, 2014). In some rare cases, such as Notch, a reporter fish line may also be available. The clear validation of our findings from phosphoproteomic analysis with some phosphospecific antibodies and the Notch-reporter line, however, indicated that phosphoproteomic analysis of embryos is a sensitive and reliable tool to estimate the activity of various signaling pathways at the whole embryo level. Due to the lack of target-specific tools, this observation is vital, since it indicates that the signaling activities of previously almost unprobeable pathways can be reliably measured with this approach.

Interestingly, our data also indicates that lapatinib and AG1478 had similar phenotypic effects, but they differentially affected signaling pathways in the phosphoproteomic analysis. Both lapatinib and AG1478 reduced the overall movement and ventricle wall growth of zebrafish embryos, but differentially perturbed the PI3K/Akt, p38 MAPK and Notch pathways. Inhibition of these differentially regulated pathways had a similar phenotypic response to lapatinib and AG1478 treatment suggesting that the differential regulation of these pathways is involved in the induced phenotypes. Interestingly, AG1478 and lapatinib-treated embryos also had differential sensitivity to Notch inhibition by DAPT. The embryos were more sensitive to Notch inhibition after lapatinib treatment and the sensitivity to Notch inhibition after lapatinib treatment has been additionally observed in the breast cancer context (Pandya et al, 2011; Abravanel et al, 2015). This further indicates that Notch signaling plays a differential role in lapatinib and AG1478-induced phenotypes. Our observations of similar phenotypes but differential signaling activities may open interesting future avenues in analyses of compounds having the same primary targets.

## Materials and Methods

### Zebrafish husbandry

Zebrafish were maintained in Aqua-Schwartz (Aqua-Schwarz, Göttingen, Germany) stand-alone units following standard procedures. The embryos were collected through natural spawning using 1.7L Sloping Breeding Tanks (Tecniplast, Buggugiate, Italy). After collection, the embryos were maintained in E3 medium in 28.5°C incubator until used in experiments. The use of zebrafish embryos under 5 days of age is not considered as animal experiment, and hence does not require ethical license according to Finnish and EU legislation. Adult zebrafish were housed under facility license MMM/465/712-93 (Ministry of Agriculture and Forestry of Finland).

### Western analysis

Zebrafish embryos of AB line at 3 dpf were treated for 1 hour with 10μM of AG1478, 30μM of Lapatinib, 30μM of Gefitinib or DMSO. After the treatment, the embryos were mechanically dissociated with a motorized pestle in 2X Laemmli buffer. The lysed samples were incubated for 5 minutes at 100°C before loading to SDS-PAGE gels. Standard protocol was used for the Western analysis. The membranes were probed with anti-phospho-p38 MAPK T180/Y182 (Cell Signaling Technologies, #9211), anti-phospho-p38 T180/Y182 (Cell Signaling Technologies, #4511) and anti-β-actin (Sigma Aldrich, A5411) primary antibodies. IR-conjugated secondary antibodies and the Li-Cor Odyssey system were used for signal detection.

### Ventricle growth assay

Zebrafish embryos of casper line at 2 dpf were treated for 2 days with 10μM of AG1478 (Cayman Chemicals, 10010244), 30μM of lapatinib (Selleckchem, S2111), 30μM of gefitinib (Cayman Chemicals, 13166), 30μM of LY294002 (Cayman Chemicals, 70920), 30μM of SB23058 (Sigma-Aldrich, 559389), 30μM of BIRB796 (Axion Medchem, HY-10320), 30μM of VT103 (MedChemExpress, HY-134955), 50μM of DAPT (Tocris, 2634) or DMSO (final concentration 1%). The embryos were anesthetized with buffered tricaine (200 mg/L) and imaged live with Zeiss AxioZoom stereomicroscope. The ventricle wall area was estimated by multiplying the average ventricle thickness with the ventricle area. The ventricle thickness and area were measured from the videos with Fiji (Schindelin et al, 2012).

### Motility assay

Zebrafish embryos of AB line at 2 dpf were treated for 2 days with 10μM of AG1478, 30μM of lapatinib, 30μM of gefitinib, 30μM of LY294002, 30μM of SB23058, 30μM of BIRB796, 30μM of VT103, 50μM of DAPT or DMSO (final concentration 1%). For the co-treatment analyses, the embryos at 2 dpf were treated for 2 days with 0.25 μM of AG1478, 0.70 μM of Lapatinib, 1μM LY294002, 0.50 μM BIRB796, 0.20 μM DAPT or DMSO alone or in the indicated combinations. The motility of the embryos was measured at 28.5°C with the Daniovision instrument (Noldus IT, Wagenigen, Netherlands) equipped with Temperature Control Unit. Tracking and analysis were performed with integrated Ethovision XT software. Outliers were removed with the method of ROUT with Q=10%.

### Notch-reporter assay

Zebrafish embryos of the tp1:Venus-pest reporter line (Ninov et al, 2012) were treated for 1 day with 10μM of AG1478, 30μM of lapatinib, 30μM of gefitinib or DMSO at 2dpf. The treated embryos were transferred into 96-well plate and imaged with Nikon Eclipse Ti-2 with 475/28nm LED excitation, Quad filter (84000v2, Chroma Technology, Olching, Germany) and 2x Nikon Plan-Apochromat (NA 0.06) objective. The intensity of the fluorescent signal was measured from the images with Fiji (Schindelin et al, 2012). The intensity values were normalized against the background signal and the background signal value was reduced from the final normalized intensity values.

### Phosphopeptide enrichment

Zebrafish embryos of the casper line at 3 dpf were treated for 1 hour with 10μM of AG1478, 30μM of lapatinib, 30μM of gefitinib or DMSO. After treatment, the embryos were mechanically dissociated with a motorized pestle in 8M Urea in 50 mM Tris-HCl, pH 8. Proteins were reduced with 10 mM D,L-dithiotreitol in 50 mM Tris-HCl, pH 8, and alkylated with 40 mM iodoacetamide in 50 mM Tris-HCl, pH 8. Samples were digested overnight with sequencing-grade modified trypsin (Promega). Peptides were desalted with Sep-Pak C18 well plate (Waters). The phosphopeptide enrichment was performed with High-Select TiO_2_ Phosphopeptide Enrichment for complex samples -kit (Thermo Fisher Scientific) according to the manufacturer’s protocol.

### Mass spectrometry

The MS/MS analysis was performed in the Proteomics Core of Turku Bioscience Centre, partially supported by Biocenter Finland. The enriched phosphopeptides were re-suspended into 0.1% formic acid before injection to LC-ESI-MS/MS protein analysis. The LC-ESI-MS/MS analyses were performed on a nanoflow HPLC system (Easy-nLC1000, Thermo Fisher Scientific) coupled to the Q Exactive HF mass spectrometer (Thermo Fisher Scientific, Bremen, Germany) equipped with a nano-electrospray ionization source. Peptides were first loaded on a trapping column and subsequently separated inline on a 15 cm C18 column (75 μm x 15 cm, ReproSil-Pur R13aq 3 μm 120 Å C18-AQ, Dr. Maisch HPLC GmbH, Ammerbuch-Entringen, Germany). The mobile phase consisted of water with 0.1% formic acid (solvent A) and acetonitrile/water (80:20 (v/v)) with 0.1% formic acid (solvent B). A 90 min 2-step gradient from 7% to 23% in 68 mins and to 36% of eluent B in 22 mins, followed by a wash stage with 100% of eluent B was used to elute peptides. MS data was acquired automatically by using Thermo Xcalibur 4.1 software (Thermo Fisher Scientific). In the DIA method, a duty cycle contained one full scan (400–1000 m/z) and 40 DIA MS/MS scans covering the mass range 400–1000 with isolation width 15 m/z.

Additionally, six gas phase fractionation (GPF) -DIA acquisitions (Searle et al, 2018) were acquired of a phosphopeptide sample pool (60,000 precursor resolution, 30,000 fragment resolution, AGC target of 1e6, max IT of 50 ms, NCE of 30) using 3 m/z precursor isolation windows with optimized window placements for phosphopeptides by EncyclopeDIA (i.e., 400.362–520.416 m/z, 500.407–620.462 m/z, 600.453–720.507 m/z, 700.498–820.553 m/z, 800.544–920.598 m/z, and 900.589–1020.644 m/z) (Pino et al, 2020).

### Mass spectrometry data analysis

Data processing was carried out by Spectronaut 14.7. software (Biognosys AG, Switzerland). A spectrum library was generated from analyzed samples and GPF-DIA acquisitions using Pulsar search engine in Spectronaut with a FASTA file of *Danio rerio*, downloaded from SwissProt (23.11.2020). Following modifications were used in the Spectronaut analysis: Carbamidomethyl C, Acetyl Protein N-term, Oxidation M, and Phospho STY. The library was used to perform DIA library -based analysis in Spectronaut to search sample raw files. The LFQ intensities of the phosphopeptides were sample-wise normalized to the sum of the LFQ intensities of all peptides and these normalized intensities were used for statistical testing. Mackskill’s test was used for differential expression analysis with the different replicate experiments assigned as covariates. The P-values were FDR-corrected. Phosphopeptides with the Q-value below 0.05 and fold change above 1.5 or below 0.66 against the DMSO control treatment were considered significantly differentially expressed.

### Bioinformatic analyses

The overrepresentation analyses were performed with the Panther Overrepresentation Test (Release 2023-10-17) (Mi et al, 2019) or gProfiler (database updated on 2024-01-25) (Kolberg et al, 2023). Reactome version 85 (Release 2023-05-25) (Gillespie et al, 2022), KEGG (Release 2024-01-22) (Kanehisa & Goto, 2000), and WikiPathways (Release 2024-01-01) (Martens et al, 2021) databases were sourced for pathway annotations. The gene set enrichment analyses were performed with GSEA v 4.3.2 (Subramanian et al, 2005) with classic analysis from ranked Q-value weighted fold-change values against the DMSO control. The c2.all.v2023.2 and c5.hpo.v2023.2 gene sets from MsigDB v2023.2.Hs (Liberzon, 2014) were used for the GSEA. The human phosphoproteomics datasets were acquired from the publications of Koch, et al., Wang et al., Imami et al., Sidhanth et al., Herman et al. and Klaeger et al. (Imami et al, 2012; Koch et al, 2016; Klaeger et al, 2017; Wang et al, 2017; Herman et al, 2022; Sidhanth et al, 2022). The overrepresented pathways with FDR below or equal to 0.10 were used to analyze the pathway overlap.

### Statistical analyses

For statistical testing, GraphPad Prism v10.1 (California, USA) and R v4.3.1 (R Core Team, 2023) were utilized. The equality of variance and normality assumptions were tested with Bartlett’s, Brown-Forsythe, Kolmogorov-Smirnov, Shapiro-Wilk, D’Agostino-Pearson, and Pearson-Omnibus tests and parametric or non-parametric testing was used accordingly. For parametric testing, one-way ANOVA, Welch’s ANOVA, and two-tailed T-test were utilized. For nonparametric testing, Kruskal Wallis ANOVA and Wilcoxon rank-sum tests were utilized. For ANOVA post-hoc analyses, either Dunn’s or Dunnet’s multicomparison tests were utilized. The correlation analyses were performed with one-tailed Pearson correlation analysis. The nonlinear relationship was estimated by fitting a quadratic curve with least squares estimation. The P-value for the R^2^ of the quadratic curve was calculated with an F-test with R v4.3.1 (R Core Team, 2023). The χ^2^ test was used to estimate the statistical significance of dataset overlap.

### Data visualization

GraphPad Prism v10.1 was primarily used for data visualization. The heatmap of differentially regulated phosphopeptides was generated with Matlab 2022b (MathWorks). Hierarchical clustering and data standardization were used for the heatmap visualization. The heatmaps for phosphoprotein and pathway overlaps were illustrated with Morpheus (Broad Institute, https://software.broadinstitute.org/). The illustrations were produced with biorender.com.

## Data availability

Mass spectrometric data is available in ProteomeXchange Consortium via the PRIDE (Perez-Riverol et al, 2022) partner repository with the accession code PXD050396. Other data is available from authors upon reasonable request.

## Conflict of Interest

The authors declare no conflict of interest.

## Funding

University of Turku (I.P., K.E., A.J.), Turku Bioscience Center of the University of Turku and Åbo Akademi (I.P., K.V., J.Ö., K.E), The Finnish Cultural Foundation (K.V.), The Finnish Foundation for Cardiovascular Research (K.V.), The Maud Kuistila Memorial Foundation (K.V.), The Turku University Foundation (K.V.).

## Acknowledgements

We thank Zebrafish Core, Proteomics Core and Cell Imaging Core of Turku Bioscience Centre, partially supported by Biocenter Finland, for the expertise and access to infrastructure. For providing reagents, we wish to thank the members of the Veli-Matti Kähäri and Kari Kurppa group. Maria Tuominen is acknowledged for skillful technical assistance.

## Author Contributions (CreDit)

Conceptualization: KV, JÖ, IP, KE

Methodology: KV, JM, IP

Validation: KV, NV, IP, AJ

Formal analysis: IP, KV, JM

Investigation: KV, IP, NV, JÖ, AJ

Data Curation: KV, JM

Writing – Original Draft: KV, IP

Writing – Review & Editing KV, IP, JÖ, NV, CS, KE, AJ, JM

Visualization: KV

Supervision: IP, CS, KE

Project administration: IP

Funding acquisition: IP, KE, CS, KV

